# The Sterile Insect Technique can efficiently reduce the reproduction of the spotted wing drosophila (*Drosophila suzukii*) in strawberry

**DOI:** 10.1101/2023.04.18.537326

**Authors:** B. Gard, A. Panel, A. Labbetoul, N. Bosshard, A. Xuereb, B. Cariou, A. Debelle, C. Oliva, S. Fellous

## Abstract

The spotted wing drosophila (SWD) *Drosophila suzukii* (Diptera: Drosophilidae) is a pest of soft fruit. Since its introduction in Europe in 2008 farmers struggle to protect their crops. The sterile insect technique (SIT) has proven efficient at controlling numerous fruit fly species and could be deployed to control *D. suzukii*. In recent years, key elements of SIT applied to *D. suzukii* have become available. However, field- and field-like experiments are scarce. In this experiment, we assayed the efficacy of a high-performance strain at reducing the reproduction of *D. suzukii* in complex, yet replicated and controlled conditions. Two ratios of sterile to fertile insects (5:1 and 1:1) using bisexual releases were compared to a control treatment with fertile, wild flies only. The presence of sterile individuals at a 5:1 ratio significantly reduced fly reproduction, measured after 5 days, by an approximate threefold factor. However, the proportion of infested fruits in the treated plots remained unaffected. The number of available berries in the cage appeared an unexpected determinant of fly infestation, suggesting undocumented density-dependent processes. The success of this assay opens the door to larger scales experiments, over several generations, and, in the near future, the field-evaluation of the efficacy of the SIT to control *D. suzukii*.

## INTRODUCTION

The sterile insect technique (SIT) enables the control of insect vectors of human and animal diseases as well as agricultural pests. Targeted agricultural pests are mainly flies and Lepidopteran (Bourtzis and Vreysen, 2021). This technique relies on the repeated releases of mass-reared and sterilized male insects. These mate with wild, fertile females that consequently produce unviable progeny. The SIT limits damage to crops through the reduction of pest population density, and even sometimes local eradication (Vreysen and Robinson, 2011). In classical SIT, sterility is achieved by exposing insects to ionizing radiation. This management method is reversible and assumed to have limited side-effect on the environment. It therefore represents a valuable tool to include in the design of agroecological, sustainable crop protection schemes (Krafsur, 1998).

*Drosophila suzukii* is an invasive pest, responsible of heavy damages on soft fruits and cherries in the invaded areas (Asplen et al., 2015). Since its introduction in 2008 in Europe, many management methods have been studied to control this pest, but few have revealed satisfactory (Tait et al., 2021). Currently, management of *D. suzukii* in strawberry relies essentially on prophylaxis, cultural control with the use of insect-proof nets, mass-trapping and the application of insecticides. Biological control is not yet sufficiently efficient because native parasitoids have limited impact on the population of *D. suzukii*, and the use of exotic parasitoids is still in an experimental phase (Lee et al., 2019).

In this context, the SIT could complement available protection strategies for soft fruits and cherry (Nikolouli et al., 2018). In the past few years, active research on SIT against *D. suzukii* has been conducted. Several key results have been obtained: mass-rearing technique (Aceituno-Medina et al., 2020; Sassù et al., 2019a), irradiation dose and stage of irradiation of the insect (Lanouette et al., 2017; Sassù et al., 2019b), transportation (Enriquez et al., 2021) have all been investigated. Furthermore, a first field study, conducted in 2022 on strawberry grown in greenhouse, confirms the potential of the technique to control *D. suzukii* (Homem et al., 2022). Unfortunately, the study was not replicated, casting doubts of the universality of its results. Further field and semi-field validations of the efficacy of the SIT applied to *D. suzukii* are hence needed. In this study, we report the results of a semi-field experiment that aimed at testing whether the release of sterile males reduces the reproduction of wild *D. suzukii* females in strawberry. We tested the effects of two ratios of sterile:fertile insects, 5:1 and 1:1. As no sex sorting method is currently available, we released both sterile male and females.

## MATERIAL AND METHODS

### Plant material

Strawberry plants (cv. Donna) were produced at CTIFL Balandran, in soilless conditions, in an insect-proof greenhouses so as to avoid contamination by other pests. Strawberry plants were planted in August 2020 (week 32). Plants were transferred to the cages in which the experiment was conducted once ripe and ripening berries were homogeneously present.

### Insect rearing

For fertile insects, we used a wild strain of *D. suzukii* collected in September 2021 in the region of Montpellier (France) and reared during two generations, on artificial diet, in control conditions (22°C, 70% RH, 14D:10N). To inoculate cages, F2 virgins were collected in the 12h post emergence, stored for 9 days before being released in the experimental units. In the first temporal bloc insects were aged 2 days instead of 9.

For sterile insects, we used a domesticated strain selected for greater sexual male performance. This strain was originally founded from adults captured in the Montpellier region in 2018 (details of the selection process cannot be disclosed due to intellectual property constraints). Insects were produced in control conditions (22°C, 70% RH, 14D:10N), on an artificial diet. In order to produced sterile adults, we extracted the pupae from the diet and sterilized them using X-ray at a dose of 150 Gy. Sterilization was carried within the 24h before the emergence of the adults from the pupae. Sterilized adults were released in the experimental cages, 9 days after sterilization. In the first temporal bloc insects were aged 2 days instead of 9.

### Greenhouse experiment

We compared two ratios of sterilized vs. wild *D. suzukii* to a control without release of sterile insects. Both sterile males and females were released. Experimental treatments were as follow: control with 10 wild males and 10 wild females without sterile insects, ratio 1:1 sterile to fertile males and females and ratio 5:1 sterile to fertile males and females (Tab. 1). All the insects used in the experiments, wild like sterilized, were unmated before release. For each experimental treatment, twenty replicates were spread over 4 temporal blocs separated by one week each (i.e. 5 replicates per bloc). Replicates were run in netted cages (150cm*70cm*70cm) placed in larger greenhouses in October and November 2021. Each cage contained fifteen strawberry plants (see above). All flies (sterile and fertile) were introduced at the same time in the experimental cage and were left to mate and lay eggs during five days. After the five days period, 15 fruits per cage were harvested, only berries ready for market were collected, leaving the green and pink ones on the plant.

The number of market-ready berries in each cage was counted in the third and fourth bloc, but is missing for the first two blocs. In the fourth bloc, the number of berries on strawberry plants was homogenized among cages -by chosing plants harboring similar numbers of berries -as we noticed large differences during the previous blocs.

**Table 1:** Number of sterile and fertile adults of *Drosophila suzukii* released in the three experimental treatments

| EXPERIMENTAL<br>TREATMENT | NUMBER OF<br>STERILE MALES OF<br>*D. SUZUKII* | NUMBER OF<br>STERILE<br>FEMALES OF *D.*<br>*SUZUKII* | NUMBER OF<br>WILD MALES OF<br>*D. SUZUKII* | NUMBER OF<br>WILD FEMALES<br>OF *D. SUZUKII* |
| --- | --- | --- | --- | --- |
| CONTROL | 0 | 0 | 10 | 10 |
| RATIO 1:1 | 10 | 10 | 10 | 10 |
| RATIO 5:1 | 50 | 50 | 10 | 10 |

### Observations

Harvested fruits were stored individually in plastic cup during 48h at 25°C and 55% RH in order to accelerate the maturation of the larvae and eggs hatching, and facilitate *D. suzukii* counts. After incubation, berries were gently scratched and dipped in a salt solution (100g/L NaCl) to induce larval exit from fruit flesh. We counted the number of *D. suzukii* eggs, larvae and pupae per fruit, all aggregated in a simple metric, offspring number in 15 berries.

### Statistical analysis

As stated above, the protocol for the first two temporal blocs was different from that of the last two. In addition to not counting the number of market-ready berries in both blocs, the flies of the first bloc (2 days post emergence) appeared too young and not sexually maturity. A preliminary survey of the data showed the number of available berries (spanning a 3 fold factor in the 3rd bloc, from 17 to 49 per cage) was a key variable that needed to be taken into account to produce good quality data (Fig. 1, 2). Variation in berry availability probably explained the abnormally low number of larvae in the control treatment of bloc 2 (Wilcoxon test of the effect of bloc among the control replicates P= 0.017). Due to these observations, we elected to only consider blocs 3 and 4 for the analysis.

**Figure 1:**
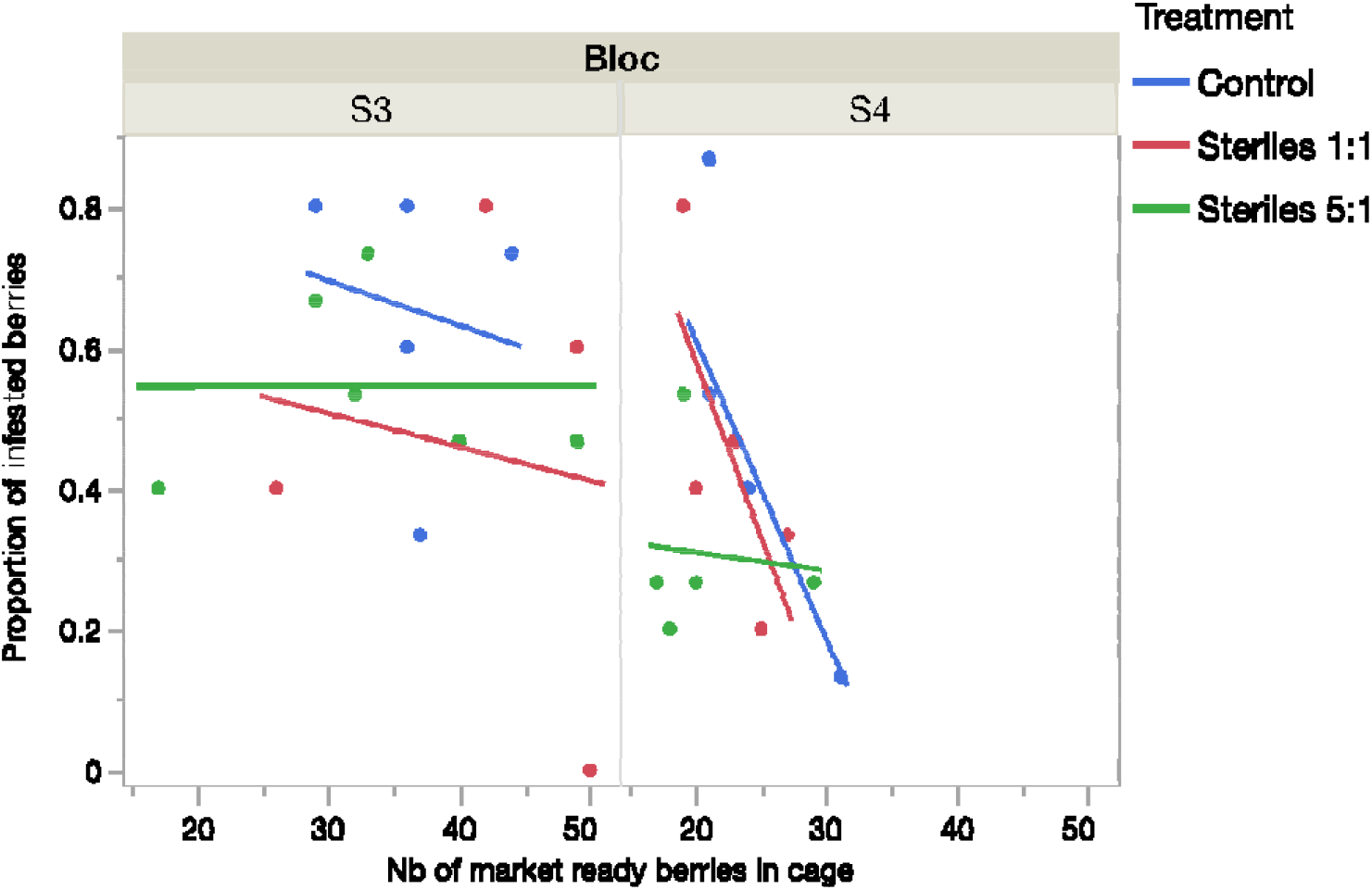
proportion of berries with at least one *D. suzukii* egg, larva or pupa.

**Figure 2:**
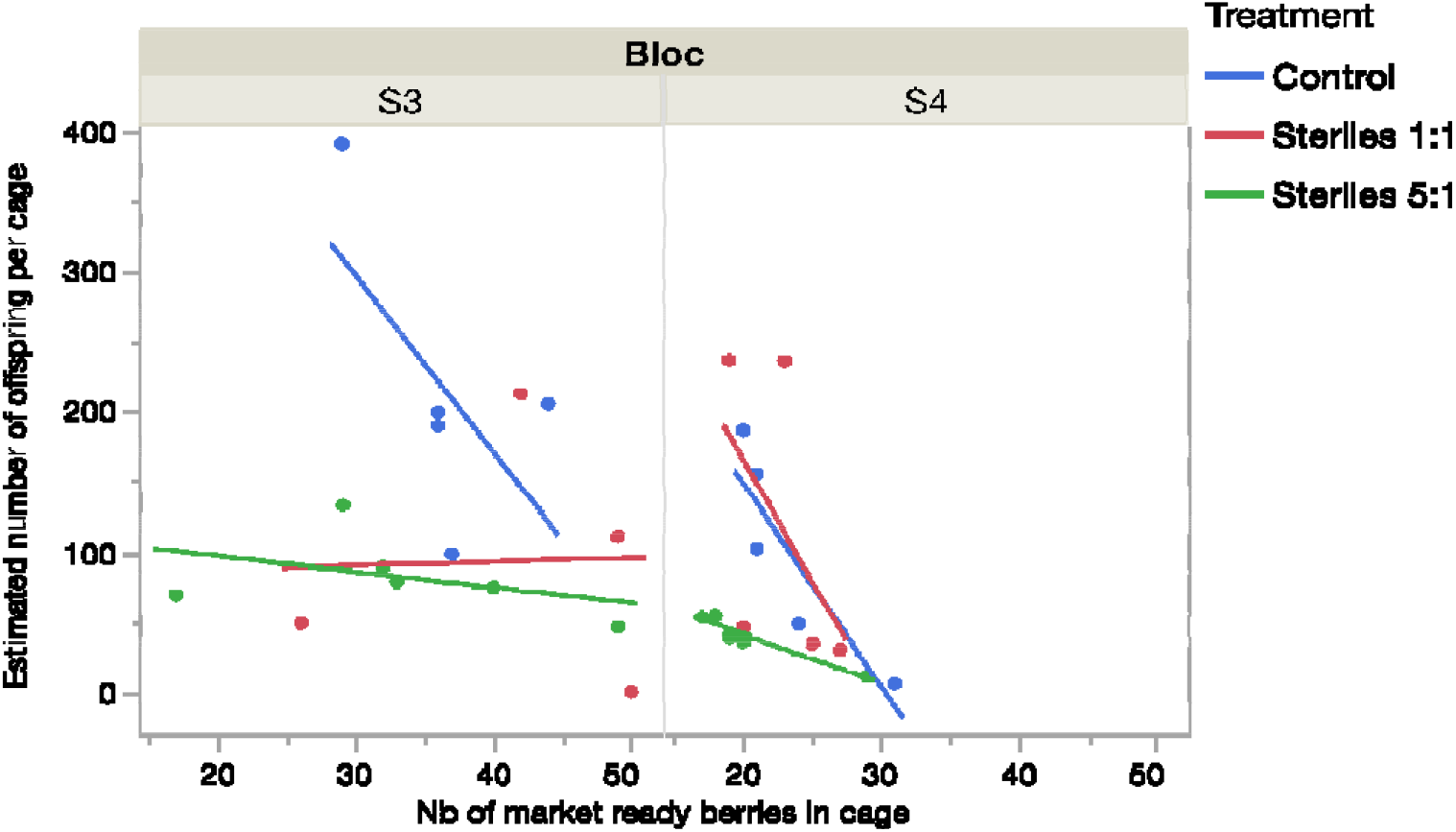
Number of *D. suzukii* offspring per cage, estimated from the number of offspring in a 15 fruits sub-sample and scaled to the total number of marketable berries counted in the cage.

Moreover, we had to correct the number of offspring in each cage by the number of berries available to the females. Indeed, because we always sampled the same number of berries in each cage (15), berry numbers variations among cages meant our subsample represented variable fractions of fly reproductive output. For all analyses, offspring number per cage was therefore scaled to the number of market-ready berries counted during harvest (i.e. estimated offspring number = Number in 15 berries /15 * number of harvested berries).

All data was analyzed with simple linear models and we used estimated cage averages as variables. We tested the effect of the SIT treatments, the number of berries per cage and the blocs on the proportion of infested fruits and the estimated number of offspring per cage (pooling eggs, larvae and pupae and correcting for berry numbers, see above).

## RESULTS

### Effect of sterile adult release on the proportion of infested berries

The proportion of infested fruits by larvae of *D. suzukii* was 0.47 (± 0.25 s.d.) in the control treatment, and 0.37 (± 0.25 s.d.) and 0.35 (± 0.20 s.d.) in the SIT treatments (1:1 and 5:1 sterile to fertile ratios, respectively). There was no significant difference between experimental treatments (F_2,25_= 1.34, P= 0.28) and no effect of berry numbers (F_1,25_= 1.76, P= 0.20). However, significant differences were observed among blocs (F_1,25_= 5.29, P= 0.03) (Fig. 2).

The estimated number of offspring per cage was significantly affected by the SIT treatment with on average 155.4 (± 106.7 s.d.) offspring in the control treatment, and 106.1 (± 96.6 s.d.) and 62.3 (± 32.3 s.d.) in the 1:1 and 5:1 SIT treatments. (Fig. 3). The SIT treatments significantly affected the number of larvae in the cages (F_2,25_= 5.18, P= 0.013). Post-hoc student’s t tests revealed a significant difference between the 5:1 and the control treatment (t= 3.19, P= 0.004). The number of available berries tended to affect the estimated number of offspring in a marginally non-significant way (F_1,25_= 3.37, P= 0.08). Significant differences were also observed among blocs (F_1,25_= 6.2, P= 0.02).

## DISCUSSION

Our study demonstrated that the release of sterile *D. suzukii* adults can significantly reduce the number of larvae present in fruit, by an approximate threefold factor. However, the proportion of fruit infested by larvae was not different between the SIT treatments and the control.

This is one of only two studies, to date, showing the SIT can reduce *D. suzukii* reproduction in field-like conditions. Recently, Homem et al. (2022) reported that sterile releases hampered *D. suzukii* population growth in a (single) large greenhouse, which was compared to two control greenhouses. This earlier work is particularly impressive for the scale of the assay and the number of insects produced and released. It however has the caveat of not being sufficiently replicated to reveal the factors that determine SIT efficacy, or failure.

Our study has strengths and weaknesses that need to be accounted for in order to put its results in perspective. First, our study has the strength of being replicated in space and time, with 10 independent cages per treatment spread over two temporal blocs. Standard statistical methods could hence be deployed ; they show, with little doubt, that the SIT is efficient at reducing *D suzukii* reproduction. This occurred even at a fairly low sterile/fertile ratio of 5, much lower than that used by Homem et al. (2022). Moreover, because the assay was run over less than 1 fly generation (5 days) we could control numerous parameters. In particular, the exact ratio of sterile to fertile flies was known and did not vary over time due to either fly reproduction or differential death between the different classes of insects. It was further possible to study the relationship between numbers of berries on plants and larvae numbers. This was an essential parameter that widely increased the precision of larvae number estimates (Fig. 1, 2). To our surprise, our dataset also suggests that the overall reproduction of *D. suzukii* can depend on the density of available fruit locally (Fig 2). Greater berry numbers in an experimental unit appeared to reduce larvae numbers, a pattern that was most visible in the control treatment.

Our study also bears weaknesses that need to be accounted for. Because we only followed one generation, it is unknown whether the short term effects we report would translate into population scale impacts over longer time scales. When changing scale, the manner strawberries are produced will have a great influence on insect multiplication. It is well established that, to date, prophylaxis, the regular removal of infested fruit, is the most efficient solution to protect strawberry and other stone fruits from *D. suzukii* damages (Lee *et al*. 2019). In normal production conditions, it is however impossible to harvest all berries, some remain, hidden in the foliage and serve as host for *D. suzukii* larvae. This phenomenon, and probably many others, was not accounted for in the present study. Along these lines, the threefold reduction of larvae we report here may be optimistic as the data from blocs 1 and 2 (which were flawed) was visually less encouraging than that of blocs 3 and 4 (used here). Further experiments will be necessary to understand inter-bloc variations and the effects of fruit density.

Ratios of sterile to fertile insects, and the release of sterile females are two important parameters in SIT programs. In our study, the greatest sterile to fertile ratio was 5:1, this is much lower than ratios usually used to control other insects (Krasfur 1998). In the case of *D. suzukii*, Homem et al. (2022) used a 100:1 sterile to fertile ratio. However, due to population growth, their actual ratio reduced during the experiment to 10:1, and eventually 1:1. Furthermore, we introduced in the experimental cages sterile females along with the sterile males. This may have reduced the efficacy of our sterile treatment as previous studies, on other fly species, demonstrated that bisexual releases were less effective than releases of males only (Caceres et al., 2002; Hendrichs et al., 1995).

It may appear surprising that the proportion of infested fruit did not respond to sterile releases. It is however mathematically logical. In the treatment with the lowest reproduction, we observed an average of 37 larvae in 15 fruits, that is 2.47 larvae per berry. Using a *Poisson* distribution, which assumes all berries can be used for oviposition, a mean of 2.47 larvae/berry implies only 8% of the berries should be larva-free. In the control treatments, we observed on average 83 larvae for 15 fruits (5.53 larvae per fruit on average). In this situation, one expects less than 1% of fruits to be larva-free. These proportions, 8% and 1% (or 92% and 99% infested) are too low and similar to detect significant changes given the modest statistical power of our study. Reducing infestation rates from 99% to 92% is also too modest to have practical consequences. We nonetheless observed lower proportions of infested berries than expected by a Poisson distribution (Fig. 3). This may be explained by the fact that all fruits may not be equally suited for fly oviposition (some are more attractive than others), and that females may prefer to lay eggs where conspecific larvae are already present (i.e. aggregative behavior) (Tait et al., 2020).

## CONCLUSION

The use of sterile insects is promising to hamper *D. suzukii* reproduction, in particular in strawberry farms under confined settings. However, a strong reduction of fly reproduction will be necessary to ensure pest-free fruits as only few flies can spread their eggs in numerous berries. As a consequence, ans as is well-known, the SIT may be most effective at preventing population growth rather than fighting damages from established fly populations.

## ACKNOWLEDGEMENTS

This study was funded by the ‘Agence Nationale pour la Recherche (ANR)’ through the SUZUKIISS:ME research project (projet ANR-21-ECOM-0002) and the ‘Interprofession de la filière des fruits et légumes frais’ (Interfel).

